# Deep learning-based predictive identification of functional subpopulations of hematopoietic stem cells and multipotent progenitors

**DOI:** 10.1101/2022.12.19.519644

**Authors:** Shen Wang, Jianzhong Han, Jingru Huang, Yuheng Shi, Yuyuan Zhou, Dongwook Kim, Md Khayrul Islam, Jane Zhou, Olga Ostrovsky, Chenchen Li, Zhaorui Lian, Yaling Liu, Jian Huang

**Affiliations:** Department of Mechanical Engineering and Mechanics, Lehigh University, Bethlehem, PA, United States; Coriell Institute for Medical Research, Camden, NJ, United States; Shanghai Key Laboratory of Medical Epigenetics, Laboratory of Cancer Epigenetics, Institutes of Biomedical Sciences, Medical College of Fudan University, Chinese Academy of Medical Sciences, Shanghai, P.R. China; Department of Bioengineering, Lehigh University, Bethlehem, PA, United States; Health & Human Biology, Brown University, RI, United States; Cooper Medical School of Rowan University, Camden, NJ, United States; Department of Medicine, Perelman School of Medicine at the University of Pennsylvania, Philadelphia, PA, United States; Temple University Lewis Katz School of Medicine, Center for Metabolic Disease Research, Philadelphia, PA, United States

## Abstract

Hematopoietic stem cells (HSCs) and multipotent progenitors (MPPs) are crucial for maintaining lifelong hematopoiesis. Developing methods to distinguish stem cells from other progenitors and evaluate stem cell functions has been a central task in stem cell research. Deep learning has been demonstrated as a powerful tool in cell image analysis and classification. In this study, we explored the possibility of using deep learning to differentiate HSCs and MPPs based on their light microscopy (DIC) images. After extensive training and validation with large image data sets, we successfully develop a three-class classifier (we named it the LSM model) that reliably differentiate long-term HSCs (LT-HSCs), short-term HSCs (ST-HSCs), and MPPs. Importantly, we demonstrated that our LSM model achieved its differentiating capability by learning the intrinsic morphological features from cell images. Furthermore, we showed that the performance of our LSM model was not affected by how these cells were identified and isolated, i.e., sorted by surface markers or intracellular GFP markers. Prospective identification of HSCs and MPPs in Evi1^GFP^ transgenic mice by LSM model suggested that the cells with the highest GFP expression were LT-HSCs, and this prediction was substantiated later by a long-term competitive reconstitution assay. Moreover, based on DIC image data sets, we also successfully built another two-class classifier that can effectively distinguish aged HSCs from young HSCs, which both express the same surface markers but are functionally different. This finding is of particular interest since it may provide a novel quick and efficient approach, obviating the need for a time-consuming transplantation experiment, to evaluate the functional states of HSCs. Together, our study provides evidence for the first time that HSCs and MPPs can be differentiated by deep learning based on cell morphology. This novel and robust deep learning-based platform will provide a basis for the future development of a new generation stem cell identification and separation system. It may also provide new insight into molecular mechanisms underlying the self-renewal feature of stem cells.

## Introduction

Hematopoietic stem cells (HSCs) and multipotent progenitors (MPPs) are important for lifelong blood production and are uniquely defined by their capacity to self-renew while contributing to the pool of differentiating cells. As HSCs differentiate, they give rise to a series of progenitor cells that undergo a gradual fate commitment to mature blood cell(Bryder, Rossi, & Weissman, 2006; Orkin & Zon, 2008). Numerous studies have defined phenotypic and functional heterogeneity within the HSC/MPP pool and have revealed the coexistence of several HSC/MPP subpopulations with distinct proliferation, self-renewal, and differentiation potentials(Morrison, Uchida, & Weissman, 1995; Seita & Weissman, 2010). Based on their self-renew capability, they can be divided into long-term (LT), and short-term (ST) HSCs and multipotent progenitors (MPPs). In the adult mouse, all HSCs/MPPs (HSPCs) are contained in the Lineage^−/low^Sca-1^+^c-Kit^+^ (LSK) fraction of bone marrow (BM) cells(Okada et al., 1992). Higher levels of HSC purity can be achieved by using signaling lymphocyte activation molecule (SLAM) family markers CD150 and CD48(Kiel et al., 2005). It has been reported that one out of every ∼2 LSK CD150^+^CD48^-^ cells possess the capability to give long-term repopulation in recipients of BM transplants. Meanwhile, ST-HSCs and MPPs can be isolated by sorting LSK/CD150^-^CD48^-^ and LSK/CD150^-^CD48^+^ cells, respectively(Kiel et al., 2005). As an alternative, HSCs can also be subdivided by CD34 and CD135 (FLT3) expression profiles. LSK/CD34^-^CD135^-^ cells are defined as LT-HSCs, whereas LSK/CD34^+^CD135^-^ as ST-HSCs and LSK/CD34^+^CD135^+^ as MPPs(Adolfsson et al., 2001). So far, there is no evidence that those three subpopulations are morphologically distinguishable under light microscope.

Of note, for better tracking of HSC activity *in vitro* and *in vivo*, several intracellular proteins, e.g., **α**-catulin and ecotropic viral integration site-1(EVI1), have been found to be expressed predominantly in murine HSCs(Acar et al., 2015; Kataoka et al., 2011). Thus, GFP expression driven by **α**-catulin or *Evi1* gene promoters has been used to identify HSCs and track their “stemness” *in vivo* or *ex vivo*(Acar et al., 2015; Kataoka et al., 2011; Y. Wang et al., 2018).

Accumulating evidence has demonstrated that the HSC aging process is accompanied by functional decline. Specifically, HSCs from aged animals (aged HSCs) manifest an increase in immunophenotypic HSC number and a decrease in regenerative capacity compared to their counterparts from young animals (young HSCs). In addition, aged HSCs tend to differentiate more to the myeloid lineage over the lymphoid lineage. Aged HSCs also demonstrate decreased homing, increased polarity, epigenetic changes, and clonal expansion(Li et al., 2020; Mejia-Ramirez & Florian, 2020; Pang et al., 2011).

Deep learning (DL) has become the state of the art for many computer vision tasks in biomedical research(Alzubaidi et al., 2021; Esteva et al., 2021). Supervised DL builds a mathematical model based on training samples with ground-truth labels. It extracts relevant biological microscopic characteristics from massive image data. The primary algorithm for DL image classification is based on the convolutional neural network (CNN). CNN is mainly composed of convolutional layers that perform a convolution with “learnable” filters. The parameters of such filters can be optimized during the learning process(Alzubaidi et al., 2021; LeCun, Bengio, & Hinton, 2015). Of note, previous studies have demonstrated that CNN is a very powerful to predict stem cell fate based on cell morphology(Buggenthin et al., 2017; Waisman et al., 2019; Zhu et al., 2021).

In our previous work, we have successfully developed a novel DL-based platform to detect rare circulating tumor cells with high accuracy(S. Wang et al., 2020). In the present study, we explored the feasibility of using DL to identify and separate different subpopulations of HSCs and MPPs based on their morphology. Experimentally, we used a large dataset of light microscopy (DIC) images of HSCs and MPPs to train our DL model, then evaluated the efficacy of its cell type prediction by testing it with validation datasets (Fig. 1). After the DL model was established, we further tested it with HSCs and MPPs that were identified and isolated by different cell surface or intracellular makers (**α**-catulin-GFP and Evi1-GFP). Remarkably, our results demonstrated for the first time that DL can extract subtle intrinsic morphological features of HSCs and MPPs from light microscopy cell images, by which the DL model can make reliable classifications. Furthermore, we also built another DL model which can effectively distinguish aged HSC from young HSC, making it a useful model to study HSC biology.

**Figure 1.**
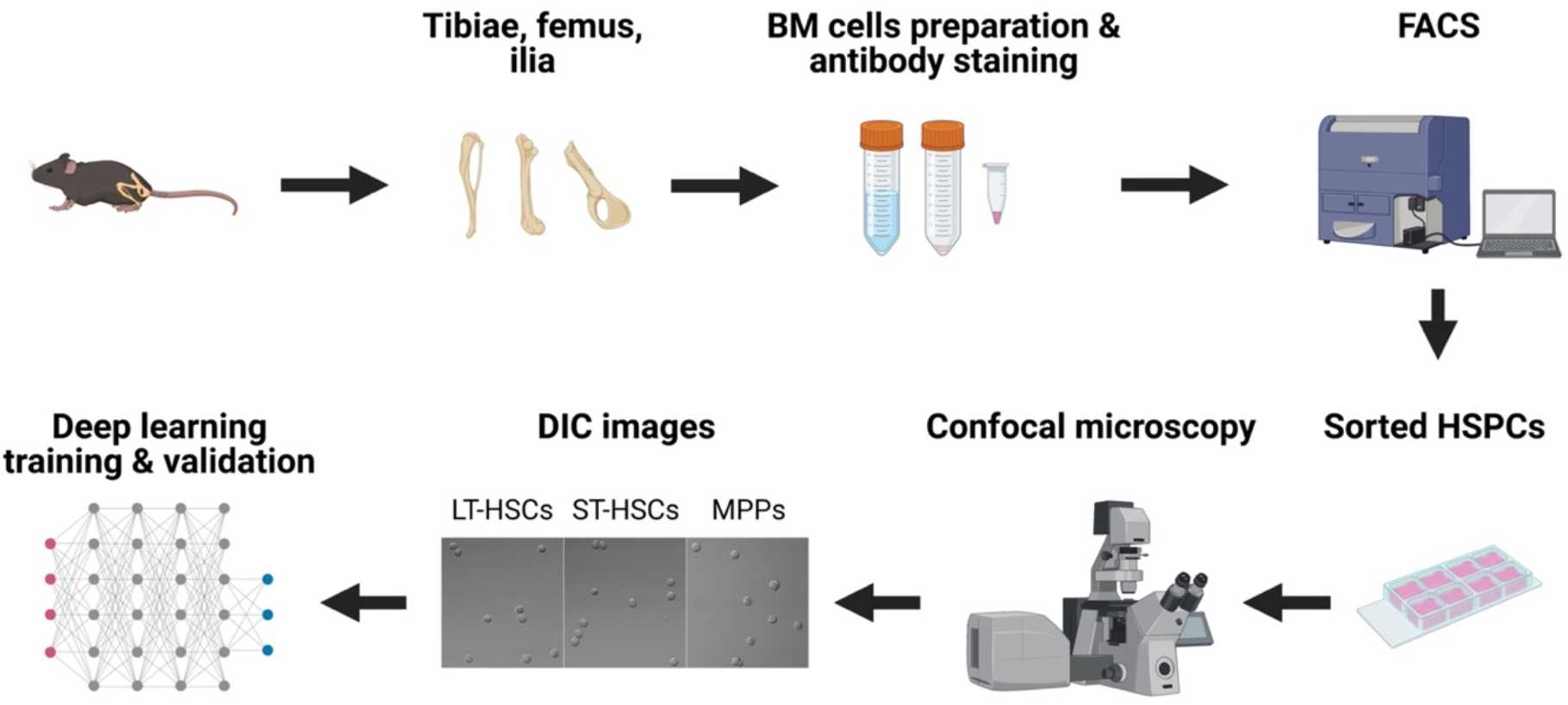
An overview of the experiment. The workflow depicts the steps from BM cell preparation, HSPCs isolation, image acquisition, to DL training and validation.

## Results

### Preparation of HSPC subpopulations and image data sets for DL training

To explore whether we can use DL to distinguish different subsets of HSPCs based on their morphology, we first sorted HSCs and MPPs from murine BM by fluorescence-activated cell sorting (FACS). We used a well-established combination of surface markers consisting of LSK (lineage^-^Sca1^+^c-Kit^+^) and SLAM (CD150 and CD48) markers and sorted out three subpopulations: LT-HSCs (LSK/CD150^+^CD48^-^), ST-HSCs (LSK/CD150^-^CD48^-^) and MPPs (LSK/CD150^-^CD48^+^) (Fig. 2A). We then seeded those cells in culture chambers with glass bottoms and acquired DIC and confocal fluorescence images (Fig. 2B). Over 96% of the recorded cells in the images exhibit anticipated fluorescence features (Fig. 2B), indicating that the sorting process was accurate and reliable. In DIC images, most HSCs (∼95%) have a spherical shape (Fig. 2B) while the rest are irregular or polymorphic. In all three cell groups, most cells appear to have a rough cell membrane, but no cell population-specific morphological features can be identified by visual inspection. In the MPP group, there tend to be more cells with diameters greater than 15 *μm* (the average size of whole BM cells is around 6.8 *μm*). However, the cell sizes of the three cell groups are mostly overlapped (Fig. 2C), making differentiation only by cell size impossible.

**Figure 2.**
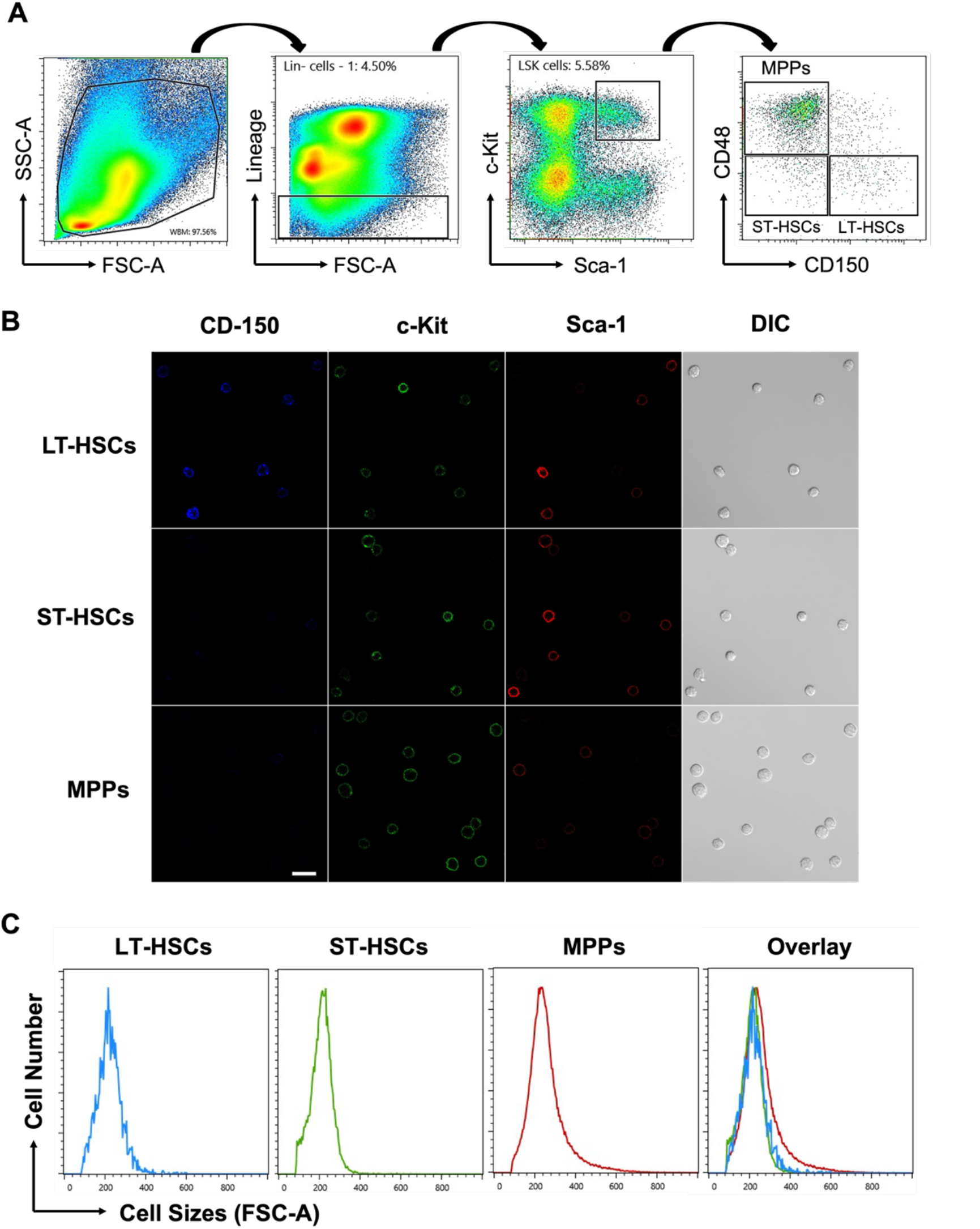
FACS of murine HSCs and MPPs using LSK/SLAM markers and cell imaging. (**A**) Representative FACS density dot plots showing the gating strategy employed to identify and isolate LT-HSCs, ST-HSCs, and MPPs from BM. (**B**) DIC and fluorescence images were taken immediately after FACS. Typical images are shown as an example. Scale bar = 10 *μm*. (**C**) Typical cell size distribution of sorted HSCs and MPPs on FACS plots.

### Development of a novel DL model to distinguish LT-HSCs, ST-HSCs, and MPPs

To build a DL model to distinguish murine HSCs and MPPs, we first took advantage of a customized MATLAB toolbox to automatically locate individual cells in acquired DIC images and segmented them into cell-centered single-cell image crops of 64 × 64 pixels. The image crops were labeled by cell types as ground truth. From 5 independent experiments, we compiled an image data set for DL model training and validation, comprising 4,050 LT-HSCs, 7,868 ST-HSCs, and 9,676 MPPs, respectively. We applied data augmentation to enhance data diversity and avoid overfitting, practiced oversampling to balance the significance of the minority LT-HSCs subset, and employed transfer learning to obviate the need for bigger data sets (detailed information described in Materials and Methods).

We designed the new DL model as a three-class classifier that would eventually assign three scores to every cell tested. The scores are between 0 and 1 with the sum of three scores being 1, representing the probabilities of being each one of the three cell types. The predicted cell type is determined by the highest score (prediction score) that ranges from 0.34 to 1 in theory. After several rounds of training and validation, the resulting DL model was challenged with cells it had never seen before. The confusion matrix in Fig. 3A summarized the results. For 647 immunophenotypic LT-HSCs tested, 60% were classified as such (consistency rate is 60%), with the rest 30% being classified as ST-HSCs and less than 10% as MPPs. Similarly, the consistency rates of classification of ST-HSCs and MPPs were 77% (1,206 / 1,574) and 77% (1,497 / 1,935), respectively. Based on these results, the performance metrics were generated to reveal various aspects of the method (Fig. 3B), including precision (positive predictive value), recall (sensitivity), macro average (arithmetic mean), and weighted average (average adjusted by sample sizes). We evaluated the performance of the model by F1 score that combines precision and recall into a single score, and the area under the curve of the receiver operating characteristic (ROC-AUC). The model has achieved an overall F1 score of 0.74 (Fig. 3B) and high AUCs on the classification of all three cell types (Fig. 3C). Together, these data suggest that DL can successfully differentiate LT-HSCs, ST-HSCs, and MPPs, and the trained model is now capable of classifying these three cell types. It will henceforth be referred as LSM model.

**Figure 3.**
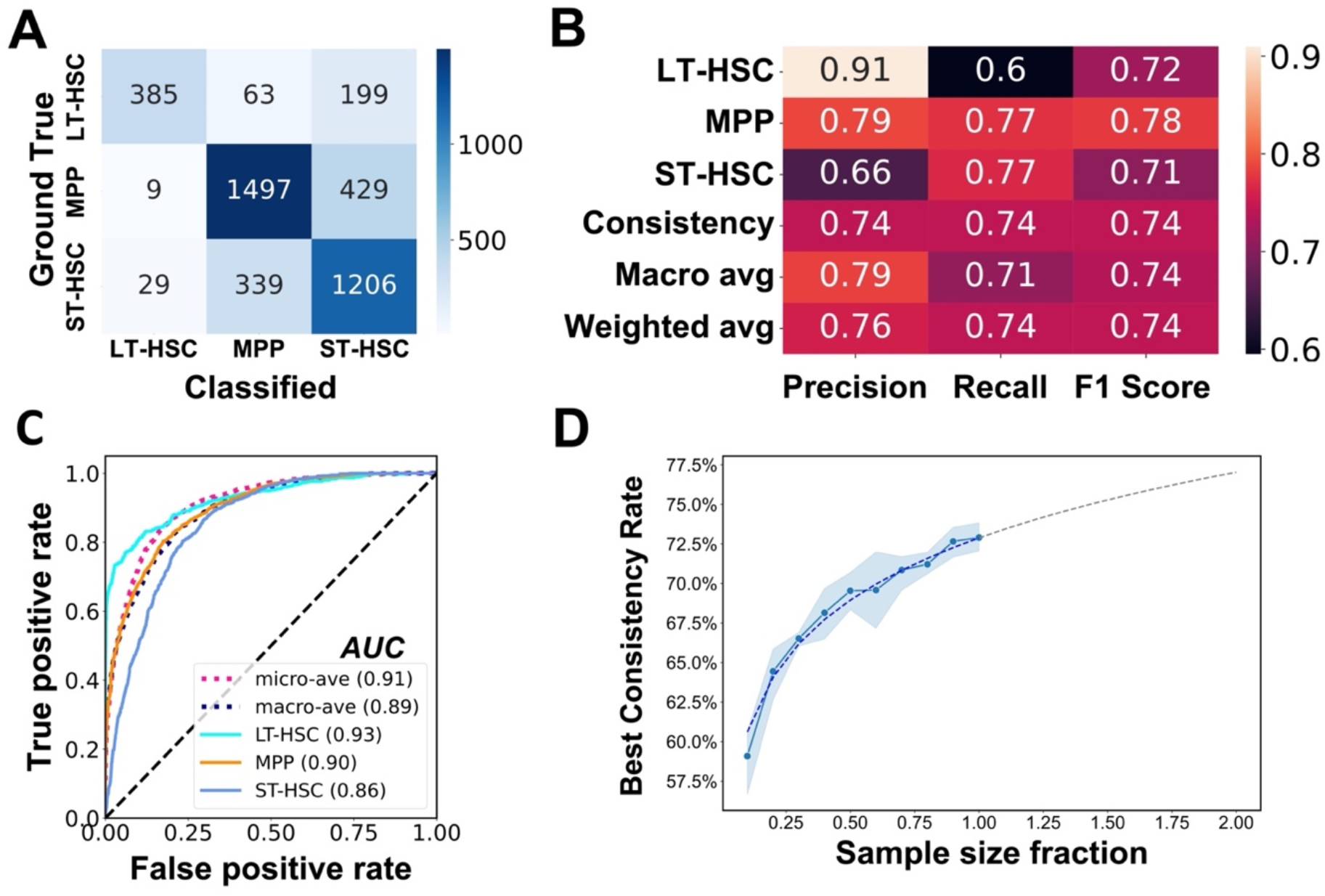
The performance of the trained DL model. (**A**) Confusion matrix of the model. After training and validation process, new cells from three HSPC subsets were classified by the model and the results are summarized in the confusion matrix. (**B**) Performance metrics of the trained model. (**C**) ROC-AUC curve of the trained model. All AUC values are above 0.85, indicative of good performance of the model. Diagonal dashed line represents a random classifier with no discrimination. (**D**) LSM model’s performance is positively correlated with the training sample size. With increased sample sizes (from 10% to 100% of total training sample), the overall consistency rate of the model improved accordingly. For each sample size, five iterations of training were executed with 20 epochs. The mean of the overall consistency rate (blue dots) and the corresponding 95% confidence interval (blue shade) are shown. The dashed line is a fitted curve with extrapolation.

Generally, more data input leads to better DL model prediction. We therefore investigated how the size of a data set would influence the performance of LSM model. To this end, we first randomly sampled 80% of the previous training data set (17,275 cells in total) to serve as the full-scale training sample (13,820 cells) and used the remainder (3,455 cells) for validation. We divided the training sample randomly into 10 fractions (1,382 cells each fraction). While keeping all other essential parameters consistent, we then trained the model with incremental sample fractions until the entire training sample was used. We practiced 5 iterations for each training sample size, and at the end of each training, validation data set was classified, and the overall consistency rate was used to gauge the performance of the model (Fig. 3D). As anticipated, the size of training sample is positively correlated with DL model performance, however, the correlation is not linear (Fig. 3D). When 80% of the training sample was used, the performance (71%) was nearly as good as that trained with entire training sample (73%). An extrapolation based on real data points indicates that predictive performance of the model can approach around 77% if the training sample size is doubled (Fig. 3D). These data indicate that LSM model has room to improve. However, an experiment as we just described may be needed to decide whether the benefit of model improvement worth the time, effort, and cost required to expand the data set.

### LSM model differentiates cells based on their morphological features

After multiple rounds of training and validation, LSM model obtained the capability to differentiate HSCs and MPPs. To elucidate what LSM model had learned from this process, we first performed a principal components analysis (PCA) on the cell images. PCA reduced the high-dimensional information from original imaging data into two-dimensional principal components (PC1 and PC2). On PCA plot, the distribution of HSCs and MPPs is dispersed and mostly mixed (Fig. 4A, left), indicating that these cell types were not distinguishable from each other at this moment. In comparison, after being processed by LSM model, imaging data were analyzed in the same way. Strikingly, cell type specific clusters were formed with limited overlap (Fig. 4A, right). These results proved that LSM model extracted features from different cell types, which are similar in one cell type but distinct among HSCs and MPPs. Next, we constructed a class activation map (Score-CAM) from the convolutional layers of LSM model (Fig. 4B). Score-CAMs are commonly used to explain how a DL model learns to classify an input image into a particular class(Huff, Weisman, & Jeraj, 2021). On the heat map, the regions received strong attention of the DL model are colored in red, while blue colored areas are ignored. When LSM model received single-cell image inputs, its strongest attention was attracted to the areas that were almost exclusively within the cell boundaries (Fig. 4B). Taken together, our data indicate that cell information critical for LSM model classification came directly from cell morphology manifested in their light microcopy images.

**Figure 4.**
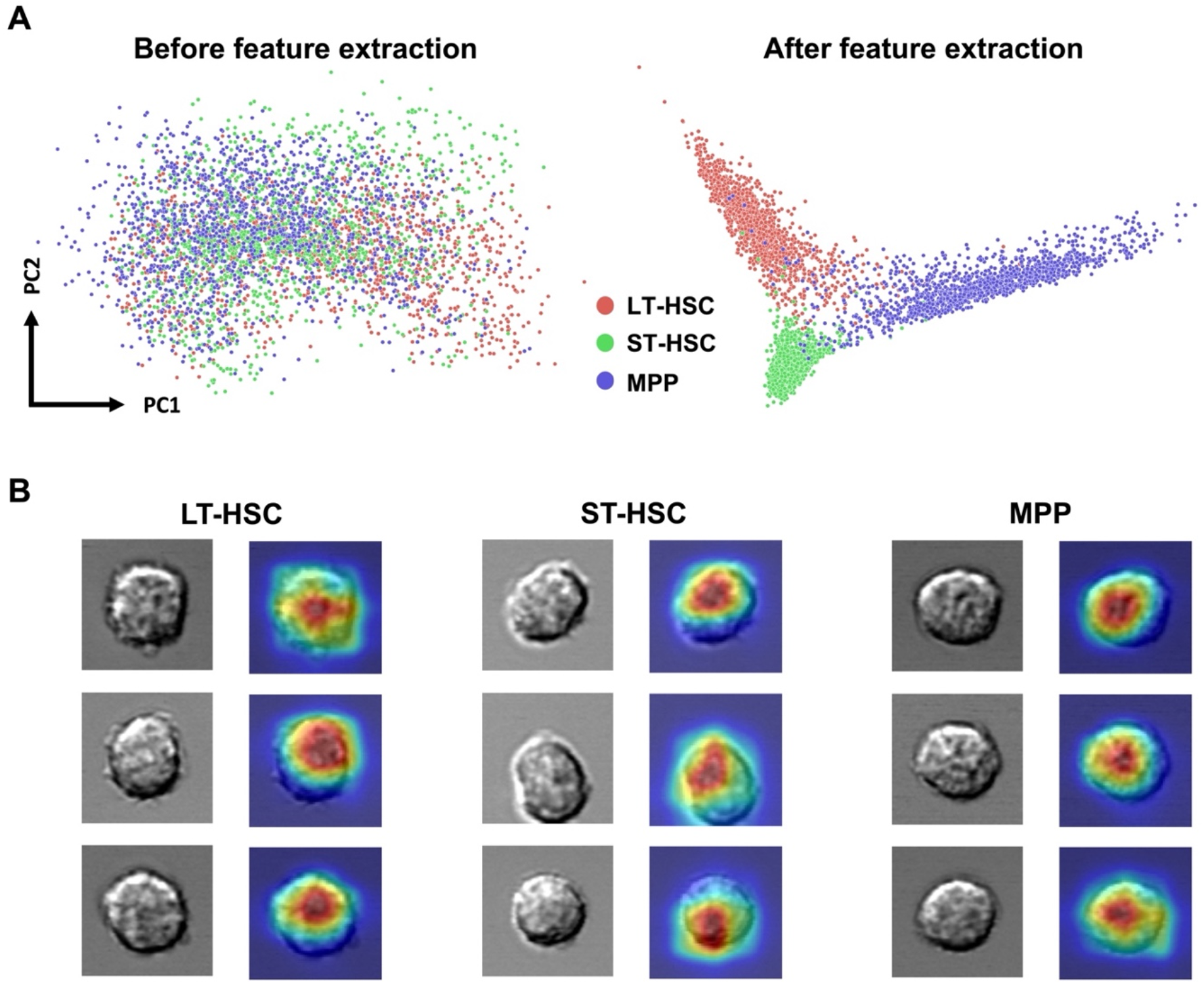
Interpretation of LSM model. **(A)** Cell imaging data were subject to a principal components analysis and the PCA score plots show the striking difference that feature extraction by LSM model can make. **(B)** Visual explanation of LSM model. Cells from the three groups were randomly selected and their attention heatmaps in LSM model were generated by Score-CAM. Red regions received the highest attention by LSM model, while blue regions were largely ignored.

### Cellular features extracted by LSM model are intrinsic to HSCs

In previous experiments, HSPCs were identified and isolated based on the binding of antibodies to corresponding surface antigens. It is possible that certain antibody-antigen interaction may result in cell-specific morphological manifestation, which makes the performance of LSM model antibody/antigen dependent. To exclude this possibility, we first applied our LSM model on HSCs and MPPs that were sorted based on LSK/CD34/CD135, another widely used set of surface markers for differentiating murine HSPCs^5^. Under this condition, the immunophenotypes of LT-HSCs, ST-HSCs, and MPPs were shown in Fig. 5A. Although the F1 scores for classification of ST-HSC and MPP are slightly lower than that seen previously, the performance of LT-HSCs classification remains consistent (Fig. 5B). These data suggest that LSM model is unlikely to capture cell morphology change induced by antibody/antigen interaction, if any, given the fact that phenotypic LT-HSCs were CD34/CD135 double negative.

**Figure 5.**
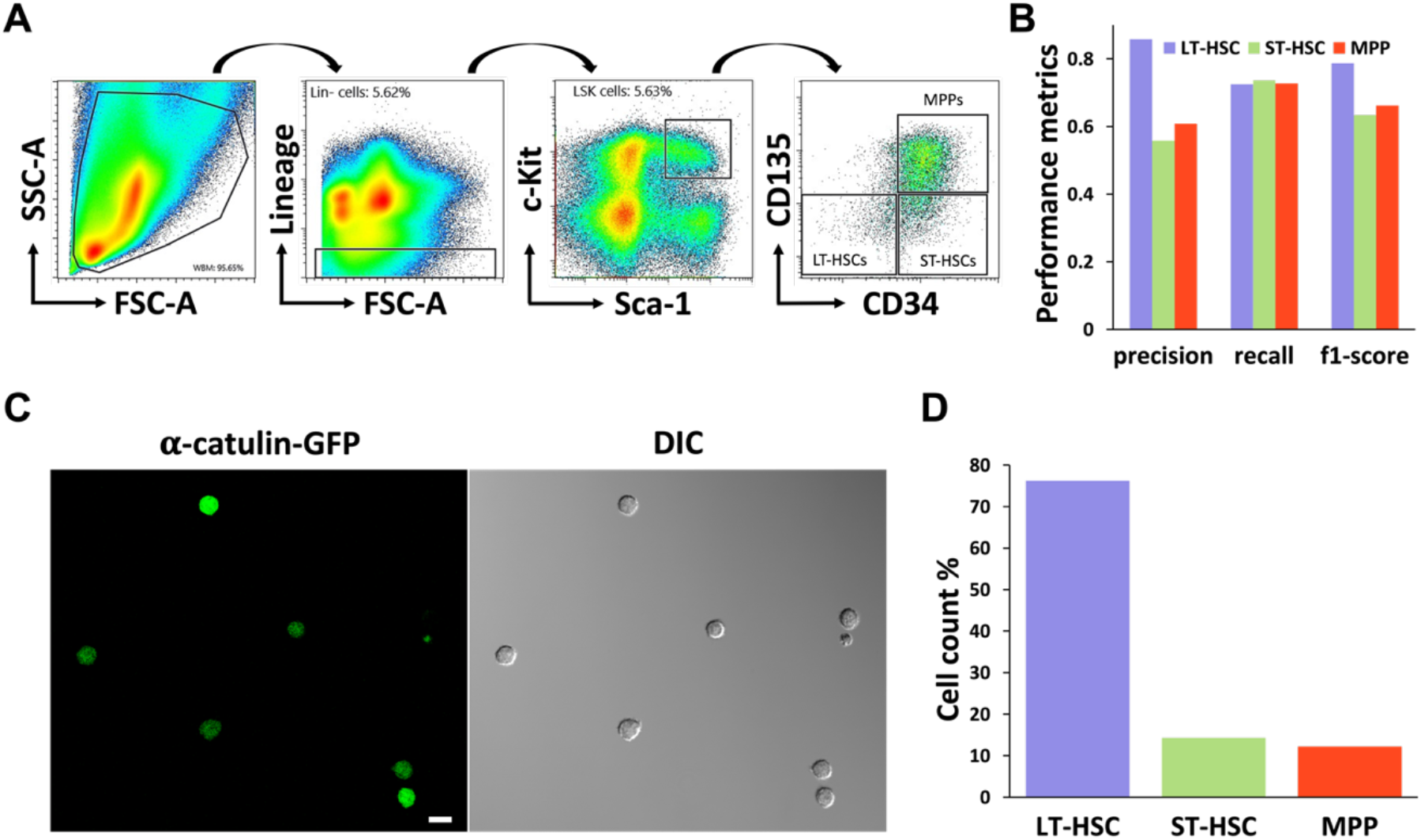
LSM model distinguishes HSPCs sorted with LSK/CD34/CD135 surface markers and **α**-catulin-GFP. **(A)** Representative FACS density dot plots show the gating strategy employed to sort murine LT-HSCs, ST-HSCs, and MPPs using LSK/CD34/CD135 surface markers. **(B)** The performance metrics of LSM model in the classification of HSPCs obtained from Fig. 5a. A total of 4,142 LT-HSCs, 873 ST-HSCs, and 1,780 MPPs were analyzed. **(C)** DIC and fluorescence images of LSK/**α**-catulin-GFP+ cells were taken immediately after FACS sorting. Representative images are shown. Scale bar = 10 *μm*. **(D)** Total 1,227 LSK/**α**-catulin-GFP+ cells were analyzed by LSM model. 74% of them were classified as LT-HSC.

We further address the issue by taking advantage of an intracellular HSC marker, the **α**-catulin protein, whose expression has been found almost exclusively in murine HSCs(Acar et al., 2015). In **α**-catulin^GFP^ mice, **α**-catulin-GFP^+^c-Kit^+^ cells in the BM are mainly LT-HSCs with a small portion of ST-HSCs(Acar et al., 2015). We therefore sorted out LSK/**α**-catulin-GFP^+^ cells and tested LSM model with acquired DIC images (Fig. 5C). A total of 1,227 LSK/**α**-catulin-GFP^+^ cells were classified. In line with expectation, 74% of them were classified as LT-HSC, 14.0% as ST-HSCs, and 12% as MPPs (Fig. 5D). Together, our data exclude the potential artificial effects of antibody/antigen interaction on cell morphology, particularly in the case of LT-HSC classification. What LSM model learned is intrinsic to the tested cells.

### LSM model can prospectively identify murine functional HSC

Our previous experiments have proved that LSM model can distinguish isolated HSPC subpopulations, we then asked how it would perform prospectively in a mixture of HSPCs when no cell-specific surface markers would be employed. To answer this interesting question, we challenged LSM model with LSK/GFP^+^ BM cells from Evi1^GFP^ transgenic mice. Evi1^GFP^ transgenic mouse is another animal model in HSC studies(Kataoka et al., 2011). Evi1 is a transcription factor of the SET/PR domain protein family and has been shown to play a critical role in maintaining HSC stemness(Kataoka et al., 2011). Unlike **α**-catulin^GFP^ transgenic mice, in Evi1^GFP^ mice where GFP reporter is controlled by Evi1 gene promoter, GFP is not only expressed in over 90% immunophenotypic LT-HSCs, 80% ST-HSC, but also in about 30% MPPs (data not shown). We tested LSM model with a data set from 1,726 LSK/Evi1-GFP^+^ cells, among which 55% were predicted as LT-HSC, 27% as ST-HSC, and 18% as MPP (Fig.6A). A fluorescence analysis revealed that GFP expression in all three predicted cell types varied greatly, however, strong GFP expression was more frequently seen in the predicted HSCs (Fig. 6B, left). In line with this finding, the average GFP fluorescence intensity of all predicted HSCs was higher than that of the predicated MPPs (Fig. 6B, left), which is consistent with previous report(Kataoka et al., 2011). This trend was not affected when the prediction score threshold (manifesting LSM model’s classification confidence) was increased to 0.5, 0.7, or 0.9 (Fig. 6B, right). Although GFP expression patterns were similar between predicted LT-HSCs and ST-HSCs, the former contained a small number of cells (6% in total tested cells) that expressed the highest level of GFP (GFP-high).

**Figure 6.**
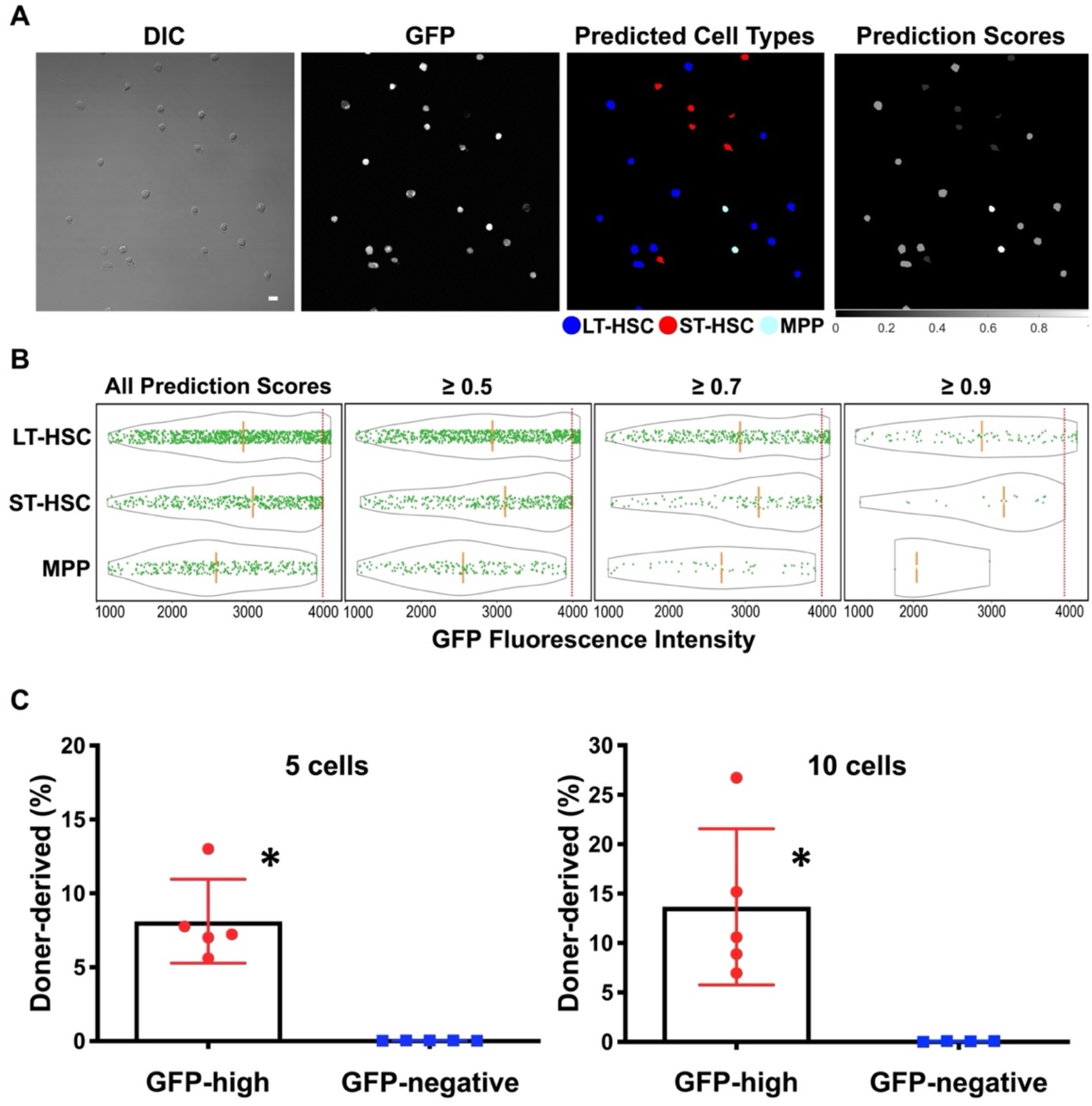
Long-term competitive reconstitution of HSCs based on prospective identification by LSM model. (**A**) LSK/Evi1-GFP^+^ cells were FACS-sorted from Evi1^GFP^ transgenic mice and cell types were predicted by LSM model. A representative field of view is shown. Scale bar = 10 ***μm***. (B) GFP fluorescence analysis in LSM model-predicted cell types. Solid green dots are individual GFP^+^ cells and the gray outlines depict the distribution of their GFP fluorescence intensity. The orange dashed lines indicate the medians of each cell groups. Increasing the prediction score threshold from 0.34 (all scores) to 0.9 didn’t significantly change fluorescence intensity features among predicted cell types, and the cells with the highest GFP expression (right to the red dotted lines) were always predicted as LT-HSC. (**C**) Competitive reconstitution of irradiated mice with GFP-high cells. Top 3% GFP expressing cells were FACS-sorted from LSK/Evi1-GFP^+^ pool and transplanted into lethally irradiated recipients. Each recipient was transplanted with 5 or 10 GFP-high cells and 3 × 10^5^ host-derived BM cells. GFP-negative LSK cells were used as control. BM was harvested after 4 months for chimerism analysis. \**P* < 0.01 (compared to GFP-negative group. Unpaired *t*-test).

It has recently been shown that in early embryonic development Evi1-high cells are predominantly localized to intra-embryonic arteries and preferentially giving rise to HSCs(Yokomizo et al., 2022). Based on this report and the prediction of LSM model, we proposed that Evi1-high precursors in adult murine BM are true functional HSCs. To test this hypothesis, we conducted a competitive transplantation experiment using FACS-sorted top 3% GFP-high cells from LSK/Evi1-GFP^+^ pool, with GFP negative LSK cells as control. We transplanted 5 or 10 GFP-high or GFP-negative cells (CD45.2) along with 3 × 10^5^ wildtype (CD45.1) “competitor” cells into lethally irradiated recipient mice (CD45.1). After 4 months, we harvested BM from the transplanted mice and measured chimerism (percentage of donor-derived cells). As shown in Table 1, the number of chimeric-positive mice — defined by convention as >1% donor-derived (CD45.2) cells in either BM or peripheral blood — was significantly higher in GFP-high group (5/5 mice for 5 cells and 5/5 for 10 cells). In contrast, no long-term reconstitution was found in GFP-negative group (0/5 and 0/4 for 5 cells and 10 cells). The degree of chimerism for GFP-high 5 cells group (mean, 8.116%) and 10 cells group (mean, 13.67%) was substantially higher than that for the GFP-negative 5 cells (mean, 0.035%) and 10 cells group (mean, 0.064%) (Fig. 6C). These results clearly demonstrated that LSM model can be used to prospectively identify functional murine HSC.

**Table 1.**
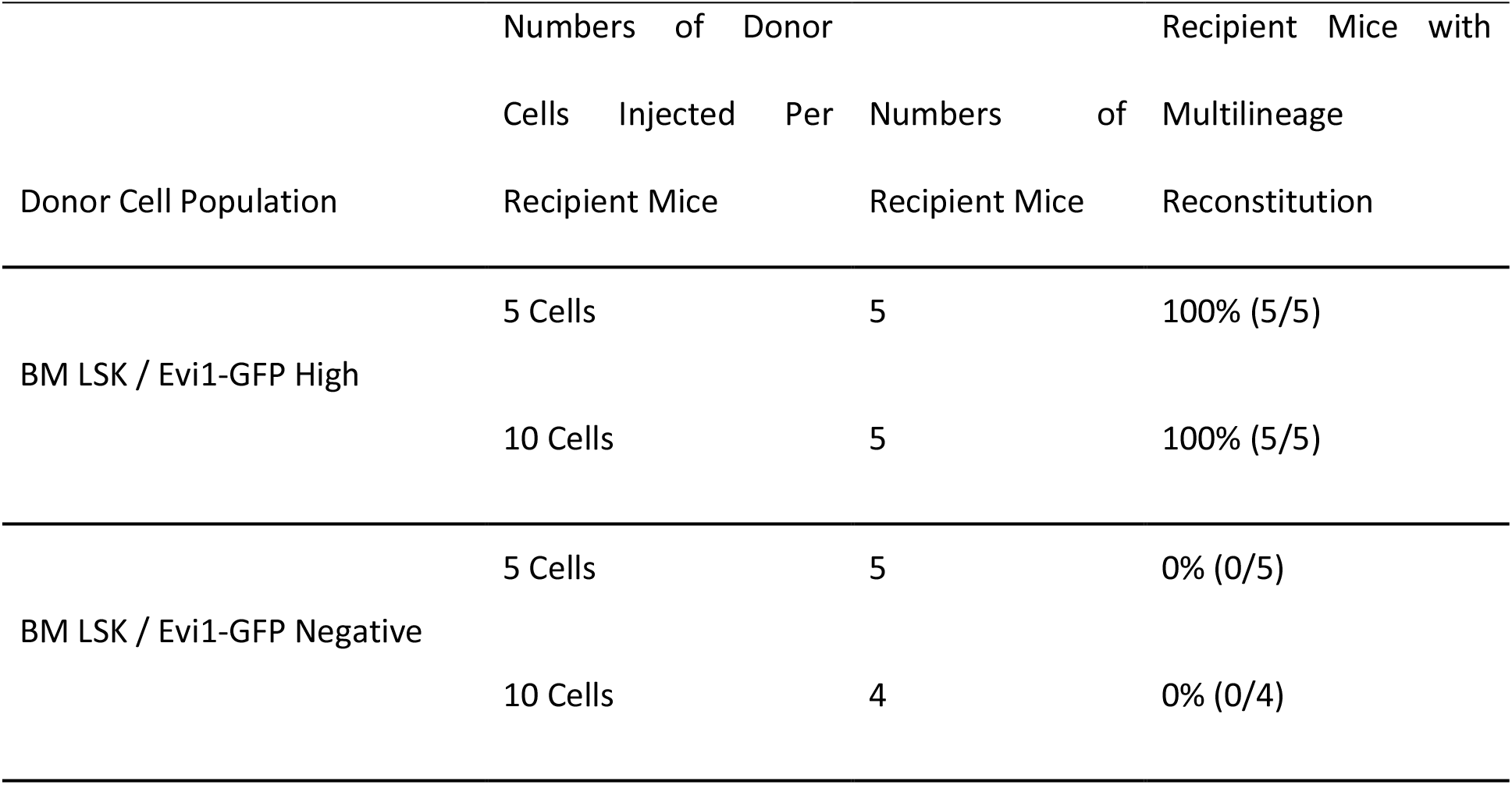
Competitive reconstitution of irradiated mice with GFP-high cells.

### Deep learning cannot differentiate MPP subpopulations (MPP2-4) from their DIC images

Accumulating evidence suggested that MPPs can be further divided into at least three subpopulations (MPP2-4), which exhibit different lineage bias and functions(Pietras et al., 2015). To test whether DL can be used to differentiate MPP subpopulations, we tried to build a new 3-class classifier exclusively for MPP classification. We adopted the same strategy for model training and validation when feeding the DL platform with DIC image data set of different MPPs (Fig. S1). After several trials, the consistency rate of classification was much lower than LSM model. Particularly, after we introduced more convolutional layers in ResNet (see Methods for details), the model performance didn’t improve, suggesting a bottleneck had been reached. By comparison, LSM model worked very well in the classification of all three MPP subpopulations (Table S1). These results indicate that while different MPP subpopulations are functionally distinct, their morphology in DIC images is largely undistinguishable. On the other hand, the data proved again that LSM model is a reliable classifier for general MPP identification.

### Deep learning can differentiate aged HSCs from young HSCs

It is well known that HSCs from aged mouse (aged HSCs) are functionally defective compared with their counterparts in young mouse (young HSCs). To investigate whether aged HSCs were morphologically different from young HSCs, we built a new DL model to distinguish aged HSCs from young HSCs. Experimentally, we first FACS-sorted LT-HSCs (LSK/CD150^+^CD48^-^) from both young (8 - 10 weeks old) and aged (24 months old) mice BM, and compiled DIC image data set. Following the same procedures in training and validation as described previously, we established a new DL model (we named it YA model) that can effectively distinguish aged HSCs from young HSCs (Figure 7). Interestingly, a small number of cells (26%) in young HSCs were classified as aged, and vice versa in aged HSCs (20% as young) (Fig. 7A). Overall, YA model identified the majority (74%) of the young HSCs as young HSCs, while classified over 80% of aged HSCs as aged HSCs (Fig. 7B). The overall F1 score of YA model is 0.78 (Fig. 7C), which is higher than LSM model. What YA model has achieved is remarkable given the fact that aged HSCs express the same set of surface markers as young HSCs. Our data demonstrated for the first time that aged HSCs are morphologically different from young HSCs, which might provide new insight into the molecular mechanism underlying the functional decline of aged HSC.

**Figure 7.**
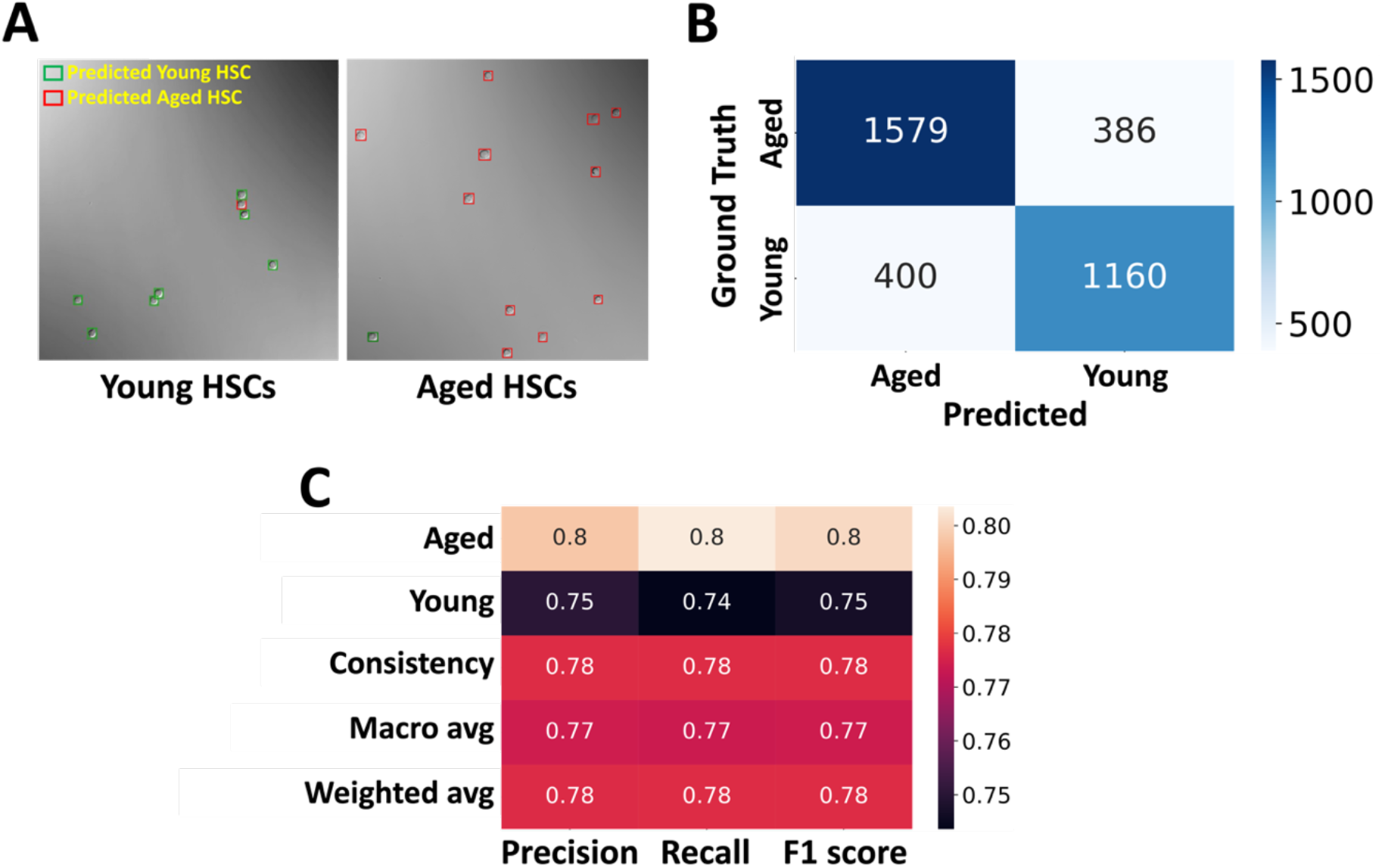
YA model differentiates young and aged LT-HSCs. (**A**) A DL model was trained to differentiate LT-HSCs from young and aged mice using image data of phenotypic LT-HSCs (LSK/SLAM/GFP^+^). Typical model predictions are shown in sample images. (**B**) After YA model was trained, it was tested with new image data set. The result was summarized in the confusion matrix. (**C**) Performance metrics of YA model.

## Discussion

Clinically, HSPCs are the most relevant component of BM transplants, which is a mainstay of life-saving therapy for patients with leukemia and congenital blood disorders. Complex combinations of cell surface markers were used to identify and separate HSPCs. In this study, we used a novel deep learning method to identify subpopulations of hematopoietic precursors. Excitingly, this new method can identify and separate subpopulations of HSPCs with excellent consistency. To our knowledge, this is the first evidence that HSPC subpopulations can be differentiated based solely on their morphology. More excitingly, we demonstrated that deep learning can effectively distinguish mouse aged HSCs from young HSCs, making it possible to identify functionally defective HSCs easily. Our novel DL models represent an important new resource for studying HSC biology. The use of these DL models in future studies should refine our understanding of stem cell morphological features and identity.

It is well known that even in the best-case scenario, the sorted HSCs and MPPs are still heterogenous. For example, less than 50% of LT-HSCs identified by SLAM markers are verified by long-term competitive reconstitution assay(Kiel et al., 2005). Similarly, the frequency of true LT-HSCs in LSK / **α**-catulin-GFP^+^ is only 33%(Acar et al., 2015). Nevertheless, we suspect that the actual frequency of true LT-HSCs might be higher because of following reasons: First, experimental procedures can affect the stemness of LT-HSCs, as evidenced by previous report that the change in environmental oxygen level greatly reduced the success rate of long-term reconstitution(Mantel et al., 2015). Second, it is possible that some of the donor LT-HSCs may not be able to engraft into the right HSC “niche” after being injected into the recipients. Third, it has been reported that the percentage of cell engraftment is higher than the percentage of long-term reconstitution(Camargo, Chambers, Drew, McNagny, & Goodell, 2006), implying that even with HSC engraftment, the condition in the niche may not be optimal for self-renewal, which makes long-term reconstitution futile. These reasons may explain why LSM model predicted a much higher percentage of cells as LT-HSC in our experiment. It is worth noting that the performance of classification can be potentially improved with more training data. However, the consistency rate appears to have a plateau even with extensive training (below 80%). At populational level, our DL models worked perfectly as they can classify different HSPC subsets accurately. However, at single cell level, the accuracy of our models is still unknown. To address this issue, long-term reconstitution assay with DL model-identified HSCs is needed.

Aged HSCs are known to be functionally defective, but they have the same immunophenotype as young HSCs(Li et al., 2020; Mejia-Ramirez & Florian, 2020; Pang et al., 2011). Therefore, it’s impossible to tell the difference between these two cell populations unless to perform a tedious long-term limiting dilution transplantation assay. However, our new YA model can accurately distinguish them based on their morphology, suggesting that YA model can be used to study HSCs with different functional states and activities.

An interesting question remains as to which morphological features are the keys for LSM model to distinguish LT-HSCs, ST-HSCs, and MPPs. It is well known that LT-HSCs, ST-HSCs, and MPPs have different self-renewal capabilities. Morphologically, how these cells are different is an important biological question meriting further study. More importantly, how aged HSCs are morphologically different from young HSCs is another key question, which warrants further investigation.

For the time being, FACS is the major method to identify and separate HSCs/MPPs. While it is a powerful analytical tool, it has several key drawbacks: it requires antibody staining and lasers as light sources to produce both scattered and fluorescent light signals. It is known that both antibody staining and intensive laser can impair cell viability and stem cell activity(Alexander et al., 2009). By contrast, DL-based platform, with further technical improvement, can be developed into an antibody-free and laser-free method to identify HSPC subpopulations. Of note, this technology may have broader applications, such as being used to identify and isolate different type of immune cells in hematopoietic system. Indeed, we also established a deep learning model which was named TB model to distinguish mouse T and B cells. Excitingly, we found that our TB model can differentiate mouse T and B cells with ∼90% accuracy (Fig. S2), indicating broad applications of deep learning technology in cell identification and separation.

### Materials and Methods Animals

C57BL/6 (CD45.2), C57Bl/6-Boy/J (CD45.1) and **α**-catulin^GFP^ mice were purchased from the Jackson Laboratory. Evi1-IRES-GFP knock-in mice (Evi1^GFP^ mice) were kindly provided by Dr. Mineo Kurokawa at the University of Tokyo ^7^. All mice were used at 8 - 12 weeks of age, except some Evi1^GFP^ mice were sacrificed at 24-month old. They were bred and maintained in the animal facility at Cooper University Health Care. All procedures and protocols were following NIH-mandated guidelines for animal welfare and were approved by the Institutional Animal Care and Use Committee (IACUC) of Cooper University Health Care.

### Antibodies

The following antibodies were used: mouse lineage cocktail-PE (BioLegend, cat# 78035), mouse lineage cocktail-APC (R&D Systems, cat# FLC001A), c-Kit-FITC (BioLegend, cat# 161603), c-Kit-APC (BioLegend, cat# 135108), c-Kit-PE/Cy7 (BioLegend, cat# 105814), c-Kit-BV421 (BioLegend, cat# 135124), Sca-1-BV605 (BioLegend, cat# 108133), Sca-1-APC (eBioscience, cat# 17-5981-82), Sca-1-BV421 (BioLegend, cat# 108127), Sca-1-PerCP-Cy5.5 (eBioscience, cat# 45-5981-82), CD150-BV421 (BioLegend, cat# 115925), CD150-PE (eBioscience, cat# 12-1501-82), CD150-PE-Cy7 (BioLegend, cat# 115913), CD48-PE/Cy7 (eBioscience, cat# 25-0481-80), CD48-BV711 (BioLegend, cat# 103439), CD48-APC-Cy7 (BioLegend, cat# 103432),CD34-FITC (eBioscience, cat# 11-0341-82), CD135-PE-Cy5 (BioLegend, cat# 135311), CD135-APC (BioLegend, cat# 135310), CD45.1-PerCP-Cy5.5 (BioLegend, cat# 110727), CD45.2-BV421 (BioLegend, cat# 109831), CD45.2-FITC(eBioscience, cat# 11-0454-82), Gr1-PE (BioLegend, cat# 108407), CD11b-APC(BioLegend cat# 101211), CD4-PE (BioLegend, cat# 116005), CD8a-PE (BioLegend, cat# 100707), B220-APC (BioLegend, cat# 103211).

### Flow cytometric analysis and cell sorting

Murine BM cells were flushed out from the long bones (tibias and femurs) and ilia with DPBS without calcium or magnesium (Corning). After lysis of red blood cells and rinse with DPBS, single-cell suspensions were stained with fluorochrome-conjugated antibodies at 4°C for 15 - 30 min. Flow cytometric analysis and cell sorting were performed on a Sony SH800Z automated cell sorter or a BD FACSAria™ III cell sorter. Negative controls for gating were set by cells without antibody staining. All data were analyzed by using either the accompanying software with the Sony sorter or FlowJo software (v.10).

### DIC image acquisition

FACS-sorted cells were plated in coverglass-bottomed chambers (Cellvis) and maintained in DPBS/2% FBS throughout image acquisition. An Olympus FV3000 confocal microscope was used to take DIC and fluorescence images simultaneously at a resolution of 2048×2048.

### Data processing

We built a MATLAB toolbox for image processing based on our previous work^23^. We used this toolbox to detect single cells in DIC images and remove the outliers (debris and cell clusters) by applying size thresholding and uniqueness checks. The toolbox then segmented the cells into cell-centered single-cell image crops of 64 × 64 pixels and labeled them by cell types. We applied data augmentation to the training examples using arbitrary image transformation including random rotation, horizontal flipping, and brightness adjustment on the original single-cell crops. We practiced oversampling on the minor classes in each run during the training experiment in order to balance different training samples. The oversampling algorithm randomly sampled training images from the minority until the number of the examples reached the same number in the majority class. Therefore, in our experiment, the training data set for each run contained equivalent numbers of data samples for all three classes.

### Deep learning framework and training

We practiced model selection with cross-validation and used ResNet-50^30^ as convolutional layers. The convolutional layers were pre-trained with ImageNet^24^ with customized starting layers to match the size of the single-cell image input, and were followed by four fully-connected layers from scratch. We applied ADAM optimizer with a weight decay of 0.05, set the learning rate of 5×10^−4^ for the fully connected layers, and fine-tuned the convolutional layers by retraining the convolutional layers at 1% of the learning rate. We reported the final training outcome with a training and validation split of 8:2 and trained the model with a batch size of 512 on a *Tesla P100 GPU* on Google Colab platform with 20 epochs via Pytorch 1.10.0.

### Long-Term Competitive Reconstitution Assays

The experiments were performed as previously described^31^. Briefly, adult congenic recipient mice (CD45.1) were lethally irradiated (1000 rad, split dose 3 hours apart). Purified donor cells were then injected along with 3×10^5^ wild type “competitor” cells (CD45.1) into the retro-orbital plexus of individual recipient mice. Hematopoietic reconstitution was monitored over time in the peripheral blood using conjugated antibodies to CD45.2 (104, FITC), B220 (6B2), Mac-1 (M1/70), CD3 (KT31.1), and Gr-1 (8C5). All recipient mice were sacrificed after 4 months and their BM cells were collected and stained with the following: Lineage cocktail-PE, CD45.1-PerCP-Cy5.5, CD150-PE-Cy7, c-Kit-APC, CD48-APC-Cy7, and CD45.2-BV421, Sca-1-BV605.

### Statistics

Statistical analysis of long-term competitive reconstitution results (Fig. 6C) was performed using an unpaired *t-*test.

## Acknowledgments

We would like to greatly thank Drs. Ying Liang at University of Kentucky College of Medicine, Guoliang Xu at Chinese Academy of Medical Sciences, Peter S. Klein at Perelman School of Medicine at the University of Pennsylvania, James Z. Wang at Pennsylvania State University, and Jean-Pierre Issa and Jaroslav Jelinek at the Coriell Institute for Medical Research for their insightful comments and discussion. We thank all the members of Liu lab and Huang lab for their help and discussions. We specially thank Steven Schneible at Coriell Institute for assistance with manuscript editing.

## Author Contributions

Y.L. and J.H. conceived and designed the project. S.W., JZ.H., Y.L., and J.H. wrote the manuscript, S.W., JZ.H. JR.H. performed the experiments with contributions from Y.S., Y.Z., D.K., MK. I., J.Z., O.O., C. L. and Z.L. S.W., JZ.H., Y.L., and J.H. analyzed and interpreted the data.

## Data and materials availability

All data needed to evaluate the conclusions in the paper are present in the paper and/or the Supplementary Materials.

