## Supplementary data for "Deep learning-based predictive identification of functional subpopulations of hematopoietic stem cells and multipotent progenitors"

### Supplementary Materials

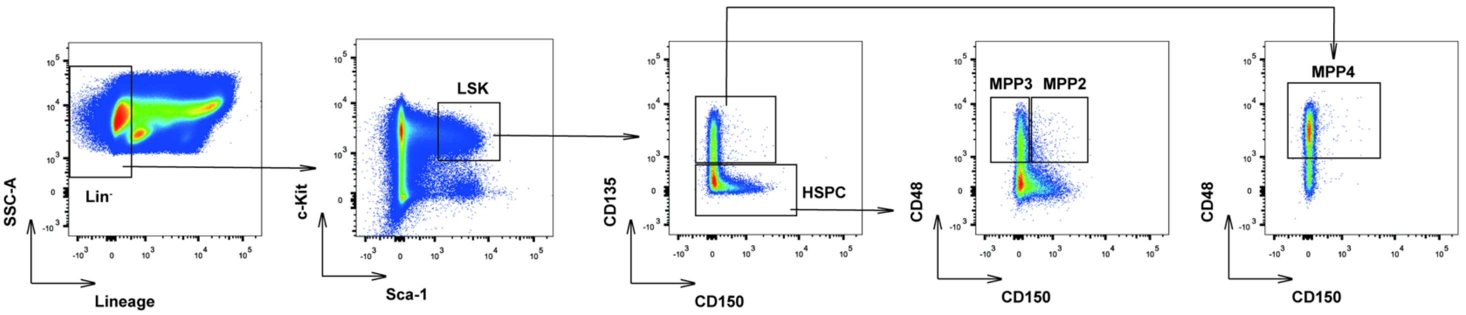


**Fig. S1. FACS sorting of different murine MPP subpopulations.** Representative FACS density dot plots show the gating strategy employed to identify and isolate MPP2, MPP3, and MPP4 from murine BM.

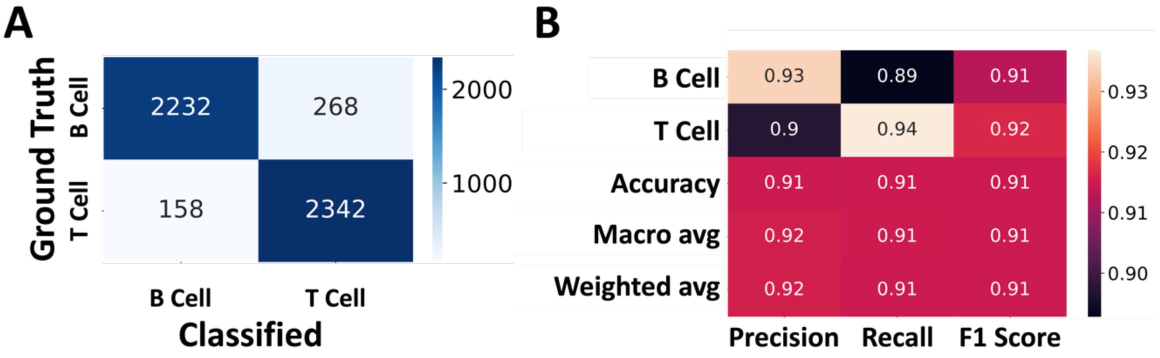


**Fig. S2. TB model differentiates T cells and B cells.** (**A**) After TB model was trained, it was tested with new image data set. The result was summarized in the confusion matrix. (**B**) Performance metrics of TB model.

**Table S1: Classification of MPP subpopulations using LSM model.**

| MPP Subpopulations | Numbers of Cells Tested | Consistency Rate of Classification Under Different Prediction Score Threshold | | |
|---|---|---|---|---|
| | | ≥ 0.34 | ≥ 0.50 | ≥ 0.70 |
| MPP2 | 477 | 90% (428/477) | 98% (392/399) | 100% (277/277) |
| MPP3 | 1490 | 90% (1346/1490) | 97% (1229/1262) | 99% (813/815) |
| MPP4 | 1301 | 90% (1170/1301) | 97% (1059/1091) | 99% (726/727) |
